# Taxonomic *vs*. Functional Diversity for the Impact Assessment of Offshore Oil & Gas Activities – An Exploratory Study on Benthic Prokaryotes

**DOI:** 10.64898/2026.07.29.741444

**Authors:** Andrea Bagi, Anders Lanzén, Jon Thomassen Hestetun, Thomas G. Dahlgren, Aud Larsen, Miriam I. Brandt

## Abstract

Improving environmental management in the offshore Oil & Gas sector requires approaches that capture the ecosystem services (ES) provided by marine sediments, particularly their roles in carbon and nutrient cycling. Microbial communities are central to these processes, and molecular tools offer new opportunities to assess their functional diversity. To explore how different sequencing approaches inform environmental impact assessment, we compared taxonomic and functional prediction based on 16S metabarcoding, shotgun metagenomics and messenger RNA-based metatranscriptomics, targeting prokaryotic communities. Our aims were to evaluate the ability of each approach to detect impact and to determine how well they captured functions relevant to ES. All approaches revealed clear differences in community composition between impacted and non-impacted sediments at both taxonomic and functional levels, with impact significantly associated with hydrocarbon and barium content. Functional inventories showed substantial overlap across the three approaches, and 48-55 ES-related processes were detectable in all datasets. While metagenomics provided the strongest statistical discrimination between impact groups, metatranscriptomics resolved the actively expressed pathways underpinning ES, yielding the most biologically meaningful functional profiles despite its lower statistical power. All approaches indicated that Oil & Gas activity drives shift towards anaerobic, hydrocarbon-degrading, and sulfur-respiring microbial communities, with hydrocarbon degradation, sulfur cycling, and metal detoxification being the dominant ES processes in impacted sediments. Metabarcoding was confirmed as a cost-effective option for impact assessment when focused on taxonomic composition. However, functional prediction from metabarcoding data proved less reliable, as several ES showed contrasting associations to impact category between metabarcoding and shotgun sequencing approaches.

## Introduction

Beyond its climate impacts, offshore Oil & Gas extraction may alter the structure and functioning of marine ecosystems, emphasizing the need for routine environmental monitoring of affected areas to support effective management [1–3]. Addressing these environmental challenges by strengthening environmental monitoring in the offshore Oil & Gas industry aligns with broader sustainability objectives reflected in the UN Sustainable Development Goals (SDGs), particularly SDG12 and SDG14, which emphasize reducing environmental impacts and protecting marine ecosystems. Incorporating an ecosystem service (ES) approach into environmental impact assessments by considering, for example, the importance of sediments in ocean carbon sequestration and biogeochemical cycling, could enhance current practices that mainly focus on macro-zoobenthic diversity and indicator species as proxies for environmental condition. Focusing more directly on ecosystem functioning and ES-relevant processes affected by disturbance may provide a more sensitive basis for detecting environmental impacts and informing mitigation needs [4].

Marine sediments provide a range of ecosystem services, *i.e.*, benefits to humans obtained from ecosystem processes, including (1) supporting services such as element cycling, nutrient cycling, primary and secondary production, (2) provisioning services such as novel genetic resources for biopharmaceuticals, and (3) regulating services such as biogeochemical regulation and detoxification [4]. Nevertheless, shifting from a taxonomy-based to an ecosystem services-based perspective in biomonitoring can be challenging since the metabolic capability of most prokaryotic species remains unknown. While both taxonomic and functional diversity can be used to infer ecosystem health and functioning, there are several arguments for shifting the focus to functional diversity [5]. Indeed, functional profiles may be less sensitive to biogeographical effects, random demographic drift, and species dispersal limitation than taxonomic profiles [5].

Molecular methods, such as eDNA metabarcoding, can enable more scalable and holistic monitoring, although challenges persist in terms of standardisation and comparability to historical biodiversity records [5]. Methodological limitations related to database coverage can be circumvented by employing taxonomy-free approaches, while challenges, such as low signal-to-noise ratios, stemming from the heterogeneous distribution of environmental DNA, underscore the potential benefits of targeting micro-benthic communities, with particular emphasis on prokaryotes [6,7]. Prokaryotes are fundamental to food webs and play key roles in carbon and nutrient cycling, with significant implications for climate regulation [8]. Their rapid growth makes them effective as early indicators of environmental change [9]. Specifically, prokaryotic microbial communities can reliably reflect sediment health under anthropogenic pressures such as urbanization, aquaculture, and offshore oil & gas activities, and prokaryote diversity has proven to be a robust, cost-effective proxy for environmental status, sometimes outperforming macro-zoobenthos [6,10–13] . Despite being historically overlooked, support is growing for their inclusion in marine monitoring programs, with successful implementation of prokaryote-based biotic indices in salmon aquaculture [14–20] and to some extent in routine monitoring of coastal and estuarine sediment quality [21].

Monitoring changes in ES requires the mapping of functional diversity. Functional diversity profiling can be achieved in multiple ways, with the most common molecular approaches being (1) shotgun sequencing of either DNA (metagenomics) or RNA (metatranscriptomics), and (2) combined use of metabarcoding and functional prediction tools. Metagenomics is the most direct way to study the functional gene content and, if done with sufficient coverage, can provide Metagenome Assembled Genomes (MAGs) of individual “species”. Metatranscriptomics, on the other hand, can offer a snapshot of the current expression (transcription) rates of functional genes. However, transcription rates may be highly dynamic, depending on short-term variations in environmental conditions, which can make the resulting data difficult to interpret since monitoring is typically carried out only periodically. Metabarcoding, due to being restricted to amplification of a single taxonomic marker (for microorganisms typically a region of the small subunit rRNA or “16S”), can better cover organisms with low abundance at lower sequencing depth than metagenomics or metatranscriptomics. However, it excludes functional genes and introduces PCR bias, depending on the universality and characteristics of the primers and target marker used [22]. Despite these limitations, Cordier (2020) has shown that functional profiles predicted from 16S metabarcoding data can be used to infer environmental status with similar accuracy compared to taxonomic profiles [10]. Suggesting that both are suitable for impact assessment. Similar results, indicating a correlation between function and taxonomy, were obtained by Abad-Recio *et al.* in a study that also applied metagenomics and GeoChip functional microarrays on the same samples from which functional inference based on 16S metabarcoding was carried out [23]. As opposed to the results of Abad-Recio *et al.*, a recent comparison between the taxonomic information obtained from metabarcoding *vs.* metagenomics found that few taxonomy-based bioindicators of benthic impact from salmon aquaculture could be identified by both approaches simultaneously and that metagenomics revealed a larger number of bioindicators [24]. However, both studies [23,24] were limited by insufficient sample size and by sequencing libraries or data obtained from several library preparations or sequencing runs with slightly differing parameters. A comprehensive comparison of metagenomics, metatranscriptomics, and metabarcoding, on the taxonomic as well as functional level, is still lacking in the context of environmental monitoring.

In this study, we employed shotgun metagenomics, mRNA-selected metatranscriptomics and 16S metabarcoding to address the following questions: (1) How well do these three approaches distinguish between impact levels, based on community structure and function? (2) How do observed impacts of Oil & Gas operations affect the structure and function of microbial communities?

## Materials And Methods

### Site Description and Sample Collection

Sediment was collected on the Norwegian continental shelf around the Veslefrikk offshore installation, which has been operational since 1989 and was decommissioned in 2022 (Fig. S1, Table S1). Sampling was performed during the routine Norwegian offshore monitoring program [25], by Akvaplan-niva in May 2022 (full reports available at [26]), with measurement of both physicochemical parameters and macrofauna biotic indices. Sediment was collected at 9 stations (5 non-impacted, 4 impacted, depth range 175-178 m). Stations with very high total hydrocarbon concentration (THC) were excluded from granulometry and morphotaxonomic analyses. [27]

Sediment for eDNA and eRNA was taken from the top 2 cm at three locations within each “chemistry” grab. For metagenomics and metatranscriptomics, sediment (10 g) was then flash-frozen in liquid nitrogen and stored at −20°C immediately after sampling. For eDNA metabarcoding, sediment (30 g) was stored at −20°C on board. All samples were shipped at - 20°C to the laboratory in Bergen, Norway, and then transferred to a −80°C freezer until further processing.

### Nucleic acid extractions

Work was performed in an over-pressured cleanroom wearing appropriate equipment for low-DNA samples (coat, mask, hairnet, disposable gloves). All work surfaces were washed with fresh 10% bleach solution and rinsed with ultrafiltered water at the beginning and end of each extraction round.

#### Standard eDNA extraction

Sediment was thawed overnight at 4°C, homogenized manually and ∼0.5 g aliquotes were prepared in triplicate PowerBead tubes (DNeasy PowerSoil Pro kit) with subsampling blanks included to monitor contamination. DNA was extracted using a semi-automated protocol combining the DNeasy PowerSoil Pro PowerBead tubes and CD1 solution (Qiagen, Hilden, Germany), bead beating (Precellys 24 homogenizer, Bertin Technologies, Montigny-le-Bretonneux, France) and a Qiagen QIAsymphony SP instrument (Qiagen, Hilden, Germany) following [27]. Extraction negative controls were included, and DNA concentrations were measured with the dsDNA HS Assay Kit on a Qubit 3.0 fluorometer (ThermoFisher Scientific).

#### Joint DNA/RNA extractions

Sediment was thawed one hour on ice. Joint RNA/DNA extractions were performed with fresh phenol/chloroform/isoamyl alcohol solution (pH 6.5-8.0, 25:24:1) using the RNeasy PowerSoil Kit combined with the accessory DNA elution kit (Qiagen, Hilden, Germany). About 5 g of sediment were used, following the manufacturer’s suggestions for wet sediments. Extraction controls were performed alongside sample extractions. All RNA extracts were DNase-treated with the TURBO-DNA-free Kit (ThermoFisher Scientific, Waltham, MA, USA) in 50 µL reactions to remove any contaminant DNA. Concentrations in final extracts were measured using a Qubit 3.0 fluorometer with the RNA HS and the dsDNA HS Assay Kits (ThermoFisher Scientific).

### Library Preparation and Sequencing

Jointly extracted eDNA and eRNA were sent to Novogene Europe (Oxford, United Kingdom) for library preparation and sequencing. Nucleic acids were processed using Novogene’s in-house protocols and the resulting fragments with 350 bp nominal insert size were sequenced on an Illumina Novaseq 6000 platform in paired-end mode, with 300 cycles (sequencing depth of 40 Gb). Metatranscriptomics libraries were prepared following ribosomal RNA depletion step (removal of rRNA by using oligos complementary to rRNAs). Like for the metagenomic libraries, PE150 strategy was used, however, a sequencing depth of 20 Gb (corresponding to 65 million reads) was chosen here.

For metabarcoding, the prokaryote 16SV4V5 region was amplified from the standard eDNA extracts with the 515F-Y/926R primer pair [28] Click or tap here to enter text. using primers with 8 nucleotides (nt) sample tags on the 5’ end, enabling nested multiplexing of PCR products as described in [29]. The triplicate extracts were pooled for PCR, as recommended by [30] and single PCR amplifications were carried out per sample, as the study aim was to resolve the core community structures [31]. Details of PCR conditions, purification, quality control and pooling steps are described in detail in the Supplementary Information. PCR-negative controls were performed along samples PCRs, and a mock community was included as positive control (ZYMObiomics microbial community DNA standard, ZYMO research, Irvine, CA, USA). Amplicon libraries were prepared by Novogene Europe using a modified PCR-free NEBNext® Ultra™ II DNA Library Prep protocol. Libraries were quantified and pooled equimolarly and sequenced on Novaseq 6000 instruments (Illumina, San Diego, CA, USA) in 250 base pairs paired-end mode, with final loading concentrations of 300-600 pM and a 40% PhiX spike-in using NovaSeq 6000 SP Reagent Kits. Raw data was demultiplexed at Novogene based on the index in the sequencing adapters. Adapters were removed, and raw data filtered by removing reads containing adapter sequences, removing reads with N > 10% and removing reads containing bases with quality scores <= 5 over 50% of the read.

### Sequence Processing and Annotation

#### Metagenomics

Bioinformatic processing was performed by the sequencing service provider (Novogene Europe) using their standard metagenomic analysis pipeline (described in detail in the Supplementary Information). Briefly, raw reads were quality filtered, assembled *de novo*, and assembled contigs were used for gene prediction. Predicted genes were dereplicated to generate a non-redundant unigene catalogue, and quality-filtered reads were mapped back to the unigenes to estimate normalized gene abundances.

#### Metatranscriptomics

Bioinformatic processing of metatranscriptomic data was performed by the sequencing service provider (Novogene Europe) using their standard analysis pipeline (described in detail in the Supplementary Information). Raw reads were quality filtered, ribosomal and transfer RNA sequences were removed, and reads from one representative replicate per station (nine samples in total) were used in the assembly process generate a non-redundant unigene reference. Quality-filtered reads from all samples were subsequently mapped to the unigene set to estimate transcript abundances.

For both metagenomics and metatranscriptomics, unigenes were taxonomically and functionally annotated by the sequencing service provider. Taxonomic annotation was performed by aligning unigenes to Novogene’s microNR database (including sequences of Bacteria, Fungi, Archaea and Viruses extracted from NCBI’s NR database, version 2018-01-02) using DIAMOND followed by lowest common ancestor (LCA) classification. Functional annotation was based on Kyoto Encyclopedia of Genes and Genomes (KEGG, version 2018-01-01) using DIAMOND best BLAST hit assignments.

#### Metabarcoding

data demultiplexing and processing was carried out using the scripts available in https://github.com/lanzen/MetaBridge. In summary, *cutadapt* v.4.5 [32] was used to demultiplex samples based on the 8 nt sample tags, and then to re-orient reads and remove primers, discarding reads with incorrect or incomplete primer sequences. Illumina reads were then denoised using DADA2 (v.1.28) to generate amplicon sequence variants (SVs), , which were taxonomically assigned using CREST v4.3.7 [33] (https://github.com/xapple/crest4) with the SSUOME1.1 database (combining SILVA SSURef v138 and EUKARYOME v1.9.3). SV tables were refined in R v.4.4 [34]by removing potential contaminants based on subsampling, extraction, and PCR blanks using the *decontam* package v.1.24.0 [35](combined frequency and prevalence-based methods, *batch* option per library). Cross-contamination was then reduced using a custom R implementation of the UNCROSS algorithm [36]. Residual cross-contamination was addressed by matching sequences with known mock species in the mock community samples. Finally, OTUs unclassified at phylum rank or assigned to eukaryotic organelles were removed.

Functional prediction was performed using Tax4Fun2 (v1.1.5) using the refined SV table and associated sequences [37] using SILVA Ref99NR as taxonomic reference database and a minimum identity to reference sequences of 97% and *normalize_pathways* set to FALSE, following the manual [38]. The output of tax4fun2 is a relative abundance table with KEGG Ortholog numbers as features.

### Ecosystem Services Relevant Functions

To enable comparisons based on functions relevant to ES, a curated list of KEGG Orthologs (KOs) was compiled using KEGG and published sources (Table S2). The selection focused on two broad ES categories: (1) supporting services like nutrient and element cycling, and (2) regulating services such as detoxification of hydrocarbons and heavy metals. Where KEGG modules existed (e.g., nitrogen cycling: M00531, M00529), they were used to link ES functions to KOs. For processes lacking modules (e.g., organic phosphorus mineralization, alkane degradation), essential genes and their KO identifiers were manually gathered. Phosphorus-related KOs were sourced from [39], alkane degradation KOs focused on key oxygenases, and metal-related genes were drawn from recent studies [40,41]. Carbon cycling KOs were selected for their roles in CO₂ fixation and methane dynamics, reflecting their impact on greenhouse gas fluxes. The resulting ES-relevant database contained 611 KOs.

### Ecological Data Analysis and Statistics

Prevalence filtering was performed at feature level (ORFs, transcripts and ASVs for metagenomics, metatranscriptomics and metabarcoding, respectively) to retain only features that were present in all 3 grab replicates at each station.

For all the statistical tests, performed in R (v4.4.0), Bray-Curtis distances calculated from Hellinger transformed data using vegan (v2.6.10), were used to quantify community dissimilarities.

Non-Metric Dimensional Scaling (NMDS) was performed using metaMDS and visualized using ggplot2 (v3.5.1). Permutational Multivariate Analysis of Variance (PERMANOVA, adonis2, 999 permutations) was used to test the effect of impact on prokaryote communities. Multivariate homogeneity of groups dispersions was evaluated with betadisper function (999 permutations).

For each approach, bar plots of mean relative abundances across impact categories showing compositional structure at family, KO and ES-relevant metabolic process level were created using ggplot2 (v3.5.1).

Similarity Percentage analysis (SIMPER) was used to identify families, KO and ES-relevant metabolic processes contributing most to the differences in community structure and function between impacted and non-impacted sites (p < 0.05). We selected the top nine contributing features from SIMPER analysis based on the values of “average” (*p* < 0.05) and plotted relative abundances of these by impact status.

Venn diagrams were created using the online InteractiVenn tool [42].

Distance-based redundancy analysis (dbRDA) was performed using the *capscale* function from the vegan package (v2.6.10) for each dataset. Chemical explanatory variables (THC, Ba, As, Cd, Cr, Cu, Hg, Pb, Zn) were centered and scaled (mean = 0, SD = 1) using the *scale* function to minimize biases due to differing value ranges. To reduce multicollinearity among chemical parameters, Pearsons’s correlation coefficient (*rho*) and variance inflation factors (VIFs, via vif.cca) were evaluated. Chromium (Cr) showed the highest number of strong pairwise correlations (*rho* > 0.7), while VIF analysis revealed extremely high values (>100) for As, Cd, and THC. Consequently, Cr, As and Cd were removed from the analysis. THC was retained due to its biological relevance in the studied context. A final model was selected using forward stepwise variable selection via the *ordiR2step* function from vegan, identifying only THC and Ba as the most parsimonious explanatory variables for community variation.

All data and code used for ecological data analysis and statistics have been deposited to Zenodo [43].

## Results

### Sequencing and annotation results

From the 27 samples collected at nine stations, metabarcoding yielded a total of 143,025 ± 37,994 quality-filtered and curated sequences per sample representing 31,891 SVs. We obtained an average of 137±3 million metagenomic and 74±9 million metatranscriptomic sequences per sample, of which 17,090,164 ORFs (83% of all ORFs detected) and 620,152 transcripts (40% of all transcripts detected) could be assigned to at least kingdom level. The prevalence filtering step (retaining only genetic features present in all three grabs per station) resulted in the loss of 86%, 17%, and 73% of prokaryotic SVs, ORFs, and transcripts, respectively.

On the Kingdom level, Bacteria represented 96.7% of SVs, 81% of ORFs, and 32% of transcripts and Archaea was found to cover only 3.3% of SVs, 1.2% of ORFs, and 0.5% of transcripts. A total of 160, 450. and 299 prokaryotic families were detected in the 27 samples by metabarcoding, metagenomics. and metatranscriptomics respectively.

Regarding functional annotation, 34% of ORFs and 23% of transcripts could be assigned to a KEGG Ortholog (KO), yielding a total of 7,316 KOs in the metagenome and 3,516 actively transcribed KOs in the metatranscriptome. In contrast, Tax4fun2 prediction utilized on average 10% of SV sequences (passing the 97% similarity-to-reference threshold) and yielded a total of 8,000 KOs for the metabarcoding data. Overall, 38% of identified KOs were shared across all three approaches (Fig. 1). More unique KOs (n=1,492) were predicted from metabarcoding data than identified in the metagenome (n=838) or metatranscriptome (n=44) data. Of the 7,316 KOs directly inferred from the metagenome, 46% (3,405 KOs) were actively transcribed and identified in the metatranscriptome and 88% (6,441 KOs) were also present in the Tax4Fun2-predicted data (Fig. 1). To enable comparisons of ES-relevant processes, analyses were subsequently restricted to our curated ES database comprising 611 KOs representing 55 ES-relevant processes in total (Table S2). All 55 ES processes were represented in the metabarcoding-predicted dataset whereas 54 were identified with metagenomics, of which 48 (89%) were actively expressed and found in metatranscriptomics data.

**Fig. 1.**
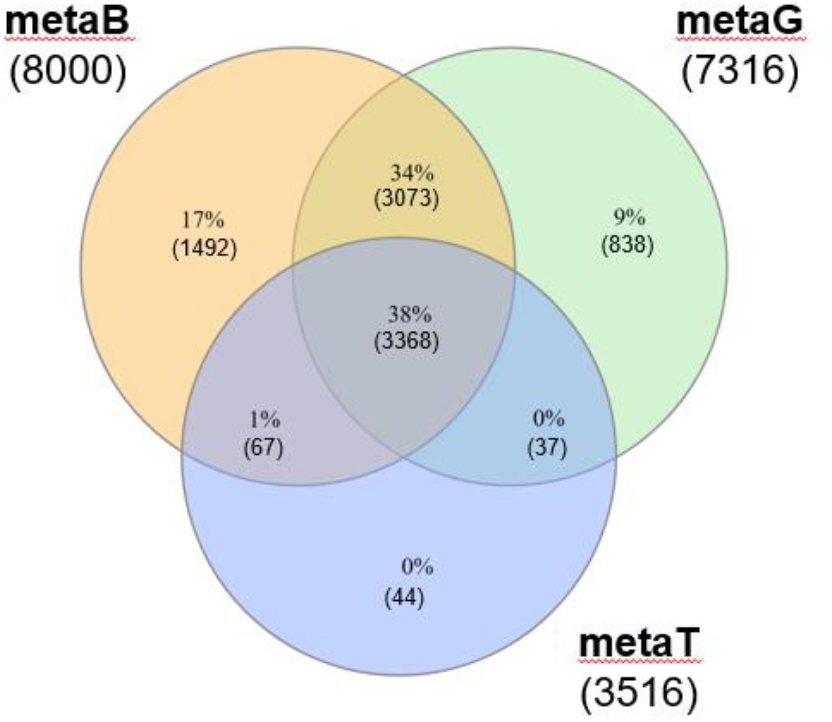
Venn diagrams of overlapping KEGG Orthologs, KOs. Overlap across all KOs identified within the three datasets. Legends include metaB = metabarcoding, metaG = metagenomics and metaT = metatranscriptomics.

### Taxa and functions affected by impact from Oil & Gas activities

Taxonomic community composition based on 16S metabarcoding data, metagenomics, and metatranscriptomics differed clearly between impacted and non-impacted sediments. At phylum level all sequencing approaches consistently showed higher relative abundances of Planctomycetota (Planctomycetes), Chloroflexi, Myxococcota, Acidobacteriota and Nitrospirae in non-impacted sediments, whereas Desulfobacterota, Bacteroidota and Campylobacterota were more abundant in impacted samples (Fig. S2). Compositions at family rank further highlighted differences between impacted and non-impacted samples (Fig. 2a). Indeed, across all sequencing approaches, Nitrosopumilaceae, Woesiaceae, Planktomycetaceae, Hyphomicrobiaceae and Rhodospirillaceae appeared to have higher abundances in non-impacted communities while Flavobacteriaceae, Pseudomondaceae, Thiotrichaceae, Desulfobacteraceae and Desulfobulbaceae dominated in impacted samples. There were a few notable differences between the family-rank profiles obtained across sequencing approaches. While dominating the metabarcoding data, Pirellulaceae, Sulfurovaceae, Desulfocapsaceae, and Sandaracinaceae were absent or less abundant in metagenomics data. Further, Rhodobacteraceae, Woesiaceae, Desulfobacteraceae, Halieaceae and Rhodospirillaceae had higher relative abundances in the metagenomes compared to the metatranscriptomic data, suggesting a relatively low metabolic activity. The opposite trend was true for Nitrosopumilaceae, Pseudomonadaceae, and Halothiobacillaceae, thus likely to be highly active.

**Fig. 2.**
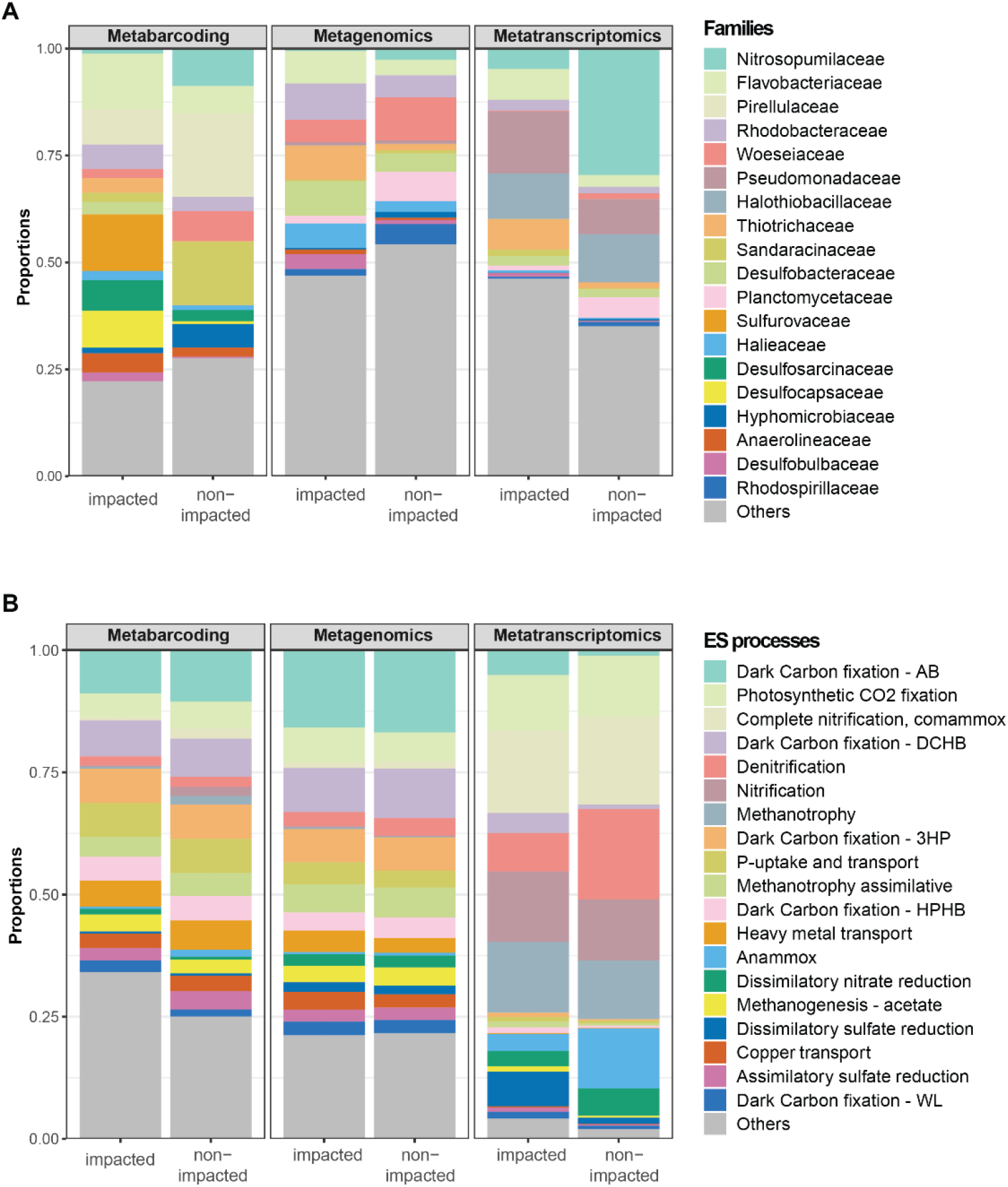
Bar plots showing relative abundances summarized by impact category for **a** family and **b** ecosystem services-relevant metabolic processes, i.e., ES processes across the three sequencing approaches.

Based on SIMPER analysis, the top ten families contributing most to differences between impact groups together explained 36% (metabarcoding), 27% (metagenomics), and 23% (metatranscriptomics) of differences in community structures between impacted and non-impacted sediments (Table S3). Considering only these top contributors, Desulfobacteraceae, Thiotrichaceae and Nitrosopumilaceae were the only families identified by SIMPER across all three approaches as significant contributors to community differences between impacted *vs*. non-impacted group. Using metagenomics and metatranscriptomics, Flavobacteriaceae was identified as associated with impacted, and Planctomycetaceae as representative of non-impacted sediments (Table S3). Other top contributing families were all unique to metabarcoding (n=7), metagenomics (n=4), and metatranscriptomics (n=4).

The taxonomic differences between impacted and non-impacted sediments were reflected in distinct functional profiles among impacted and non-impacted stations (Fig 2.b). Metatranscriptomic data showed the most pronounced differences in ES process composition between impacted and non-impacted samples (Fig. 2b). Transcripts involved in ES processes such as Photosynthetic CO_2_ fixation (i.e., Calvin-Benson cycle processes), complete ammonia oxidation (COMAMMOX), denitrification, nitrification, methanotrophy, anaerobic ammonia oxidation (ANAMMOX), dissimilatory nitrate reduction (DNR) and dissimilatory sulphate reduction (DSR) dominated the metatranscriptome. Dark carbon fixation pathways, methanogenesis, methanotrophy and DSR were dominating impacted samples, while denitrification, ANAMMOX, and nitrate reduction were considerably more abundant in non-impacted samples. None of these ES processes showed any stark differences between impact categories in the DNA-based data (Fig. 2b). ES process composition agreed largely between the metabarcoding-predicted profiles and those obtained from shotgun metagenomes, albeit some functions, *e.g.*, nitrification, metanotrophy, and ANAMMOX resulted in higher relative abundances in the Tax4Fun2-predicted profiles (Fig. 2b).

SIMPER analysis on the functional level confirmed patterns seen in the functional profiles. Indeed, metatranscriptomics better captured functional variation between impacted and non-impacted sediments (Table S3). Between group variation in the metatranscriptome was concentrated in relatively few ES-relevant processes, as only 6 of the 39 significant ES processes (15%) explained 50% of the cumulative dissimilarity between impacted and non-impacted sites, whereas metagenomics and metabarcoding required 13 of 49 (27%) and 12 of 48 (25%) significant ES processes, respectively. Further, SIMPER analysis of ES-relevant processes highlighted that non-impacted sediments were characterized by microbial communities supporting complementary nitrogen cycling pathways, including nitrification, comammox, anammox, and nitrogen fixation, together with diverse carbon turnover processes such as polysaccharide degradation, methane cycling, fatty acid degradation, and anaerobic fermentation (Table S3). The same analysis showed that impacted sediments were characterized by sulfur oxidation and sulfate reduction, together with anaerobic hydrocarbon degradation, and metal detoxification related ES processes (Table S3).

Some significant ES processes identified by SIMPER, for example ANAMMOX, appeared as a significant contributor to non-impacted sediments in both metabarcoding and metatranscriptomics data (Fig. S3). However, other significant ES processes showed contrasting patterns among sequencing approaches. For example, while nitrification and methanotrophy were representative of impacted sediments with metabarcoding, they were representative of non-impacted sediments with metatranscriptomics. Similarly, methanogenesis from amines, the single common ES process identified by SIMPER in both DNA-based methods, was indicative of impact with metabarcoding and non-impact with metagenomics. When contrasting the two DNA-based methods, predicted functional profiles from metabarcoding and directly inferred profiles from metagenomics showed a 29% overlap in ES-relevant processes identified as major contributors to impacted communities (Fig. 3a) and 26% overlap for in non-impacted communities (Fig. 3b).

**Fig. 3.**
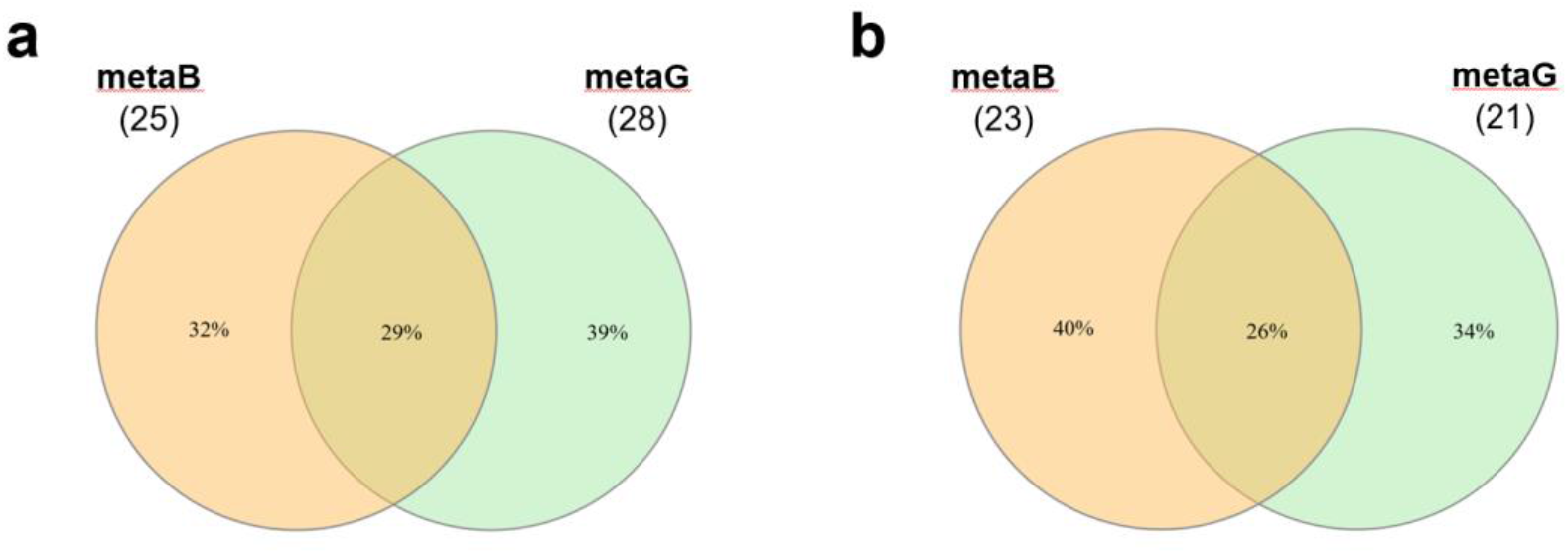
Venn diagrams of ecosystem service relevant functions (ES processes) displaying the percentage overlap between ES processes identified as major contributors in **a** impacted and **b** non-impacted communities, identified by metabarcoding and metagenomics.

### Comparative discriminative performance of the three approaches

NMDS analyses showed similar grouping of impacted *vs.* non-impacted stations across the three sequencing approaches (metabarcoding, metagenomics, and metatranscriptomics) and feature types (families, KOs, and ES-processes; Fig. S4). PERMANOVA also indicated that differences between impacted *vs*. non-impacted samples were significant for all three approaches and feature types (p < 0.001; Table 1). However, group dispersions were also significantly different (betadisper, p < 0.01; Table 1) for all tested combinations of approaches and features, with impacted samples showing much greater variability than non-impacted samples (Fig. S4).

**Table 1.** Percent explained variation in community composition attributable to impact status (impacted vs. non-impacted) based on PERMANOVA (pseudo R^2^) or a constrained ordination model corrected for the number of explanatory variables included in the model (CAPscale – adjusted R^2^)

| | PERMANOVA - pseudo $R^2$ | | |
| --- | --- | --- | --- |
|  | Families | KEGG Orthologs | ES process |
| **metabarcoding** | 0.60 | 0.55 | 0.60 |
| **metagenomics** | 0.64 | 0.64 | 0.65 |
| **metatranscriptomics** | 0.43 | 0.40 | 0.46 |

| | CAPscale - adjusted $R^2$ | | |
| --- | --- | --- | --- |
|  | Families | KEGG Orthologs | ES process |
| **metabarcoding** | 0.73 | 0.66 | 0.67 |
| **metagenomics** | 0.77 | 0.76 | 0.67 |
| **metatranscriptomics** | 0.55 | 0.53 | 0.48 |

Variation in community composition was significantly explained by THC and Ba based on dbRDA. The first constrained axis explained between 87% and 96% of the variance accounted for by these two explanatory variables across approaches and feature types, indicating that chemical load was the dominant gradient shaping community structure (Fig. 4). Metagenomics consistently explained higher proportions of compositional variation between impact statuses (Table 1) while metatranscriptomics performed poorest overall.

**Fig. 4.**
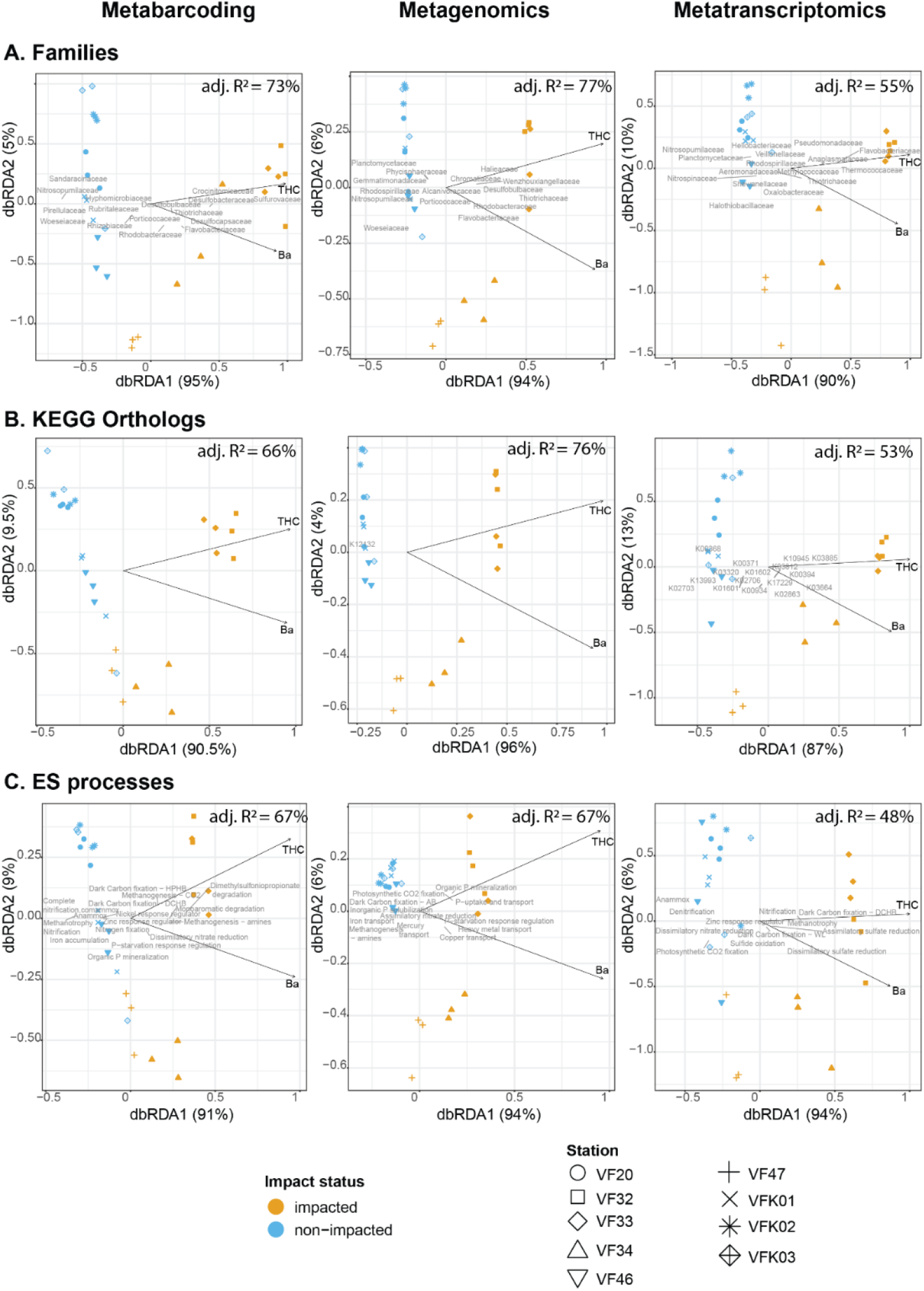
dbRDA biplots with adjusted R^2^ values of explained constrained variation for the final model that included total hydrocarbon concentration (THC) and barium (Ba) as explanatory variables. All the overall model *p*-values and per predictor *p*-values were < 0.05, except for Ba with ecosystem service relevant (ES) processes determined from metatranscriptomics, where *p* = 0.17. Panel **a** contains figures for the feature level: family, **b** for feature level: KEGG ortholog and **c** for feature level: ES processes across metabarcoding (**left**), metagenomics (**middle**) and metatranscriptomics (**right**) datasets.

According to both PERMANOVA and CAPscale, taxonomic community structure (family level) generally explained a higher proportion of the variation between impacted and non-impacted samples than functional community structure (KO or ES processes) across all three sequencing approaches. This advantage was less pronounced with PERMANOVA, where ES processes derived from metagenomics and metatranscriptomics explained more of the community variation associated with impact than taxonomic composition (Table 1). KO profiles discriminated status levels equally well as families when either metagenomics or metatranscriptomics were employed for functional profiling, while KOs performed poorer when function was predicted from metabarcoding data.

## Discussion

Our study evaluated the applicability of 16S rRNA gene metabarcoding, shotgun metagenomics, and mRNA-targeted metatranscriptomics for ecosystem service–based monitoring of marine benthic environments, both for targeting taxonomic and functional composition of prokaryotes (from metabarcoding data, function being predicted using Tax4Fun2). In general, all three approaches could explain large proportions (40-77%) of the variation between impacted and non-impacted sediments from offshore Oil & Gas installations, regardless of the type of features considered (families, KOs or ES processes). However, the taxa, functional groups, and ecosystem processes that differed depending on impact showed relatively little overlap between these three approaches, highlighting both the ability of them to provide complementary insights, and the inconsistencies or biases affecting them.

### Oil & Gas activity drives shift towards anaerobic, hydrocarbon-degrading, and sulfur-respiring microbial communities

Non-impacted sediments were characterized by an assemblage of prokaryotic families that collectively support key biogeochemical processes under both oxic and anoxic conditions for nitrogen cycling (nitrification, comammox, anammox and nitrogen fixation) and diverse carbon turnover pathways (carbon fixation, polysaccharide degradation, methylotrophy, methane oxidation, fatty acid degradation, and anaerobic fermentation). Such functional complementarity indicates a metabolically versatile community with potential to sustain multiple ecosystem services, promoting ecological stability and functional resilience. The lower transcriptional activity of the otherwise highly abundant Woesiaceae, previously reported as indicators of good environmental status [44], demonstrated that taxonomic abundance alone may not be a reliable proxy for ecosystem functioning, highlighting the value of integrating metagenomic and metatranscriptomic data.

In contrast, impacted sediments were characterised by families involved in sulphur cycling, with some also potentially participating in anaerobic and to a lesser extent aerobic hydrocarbon degradation. The dominance of sulphur oxidizing families, Sulfurovaceae (metabarcoding) and Thiotrichaceae (metagenomics and metatranscriptomics), together with the high abundance of sulphate-reducing taxa including Desulfocapsaceae (metabarcoding), Desulfobacteraceae (metabarcoding, metagenomics), Desulfobulbaceae (metabarcoding, metagenomics, metatranscriptomics) and Crocinitomicaceae (metabarcoding), all abundantly detected at impacted stations, indicates strong redox stratification within the upper 2 cm of sediment. The high activity of Thermococcaceae observed in the metatranscriptome, further supports the prevalence of anaerobic conditions at impacted stations [45,46]. Elevated abundance and activity of Desulfobulbaceae at impacted stations, together with the well-established hydrocarbon-degrading capabilities of Desulfobacteraceae suggests that anaerobic hydrocarbon degradation is currently a major detoxification pathway under these conditions [47,48]. More broadly, the dominance of Desulfobacterota also points to an important role in mercury cycling and interactions with carbon-cycling microorganisms in anoxic marine sediments [40]. These taxonomic patterns are consistent with the enrichment of ES-related functions in impacted sediments, particularly those associated with hydrocarbon degradation, sulfur cycling, and metal detoxification, linking shifts in community composition to changes in ecosystem functioning.

Overall, the microbial assemblages detected in disturbed sediments at offshore Oil & Gas installations are likely derived from natural “source” communities, both free living and symbiotic, that have evolved under analogous geochemical conditions. Comparable microbial consortia are well documented from pockmarks and other natural hydrocarbon seeps, as well as from large organic falls such as whale carcasses [49–53]. These environments share key features such as localized hydrocarbon flux, sulphidic and anoxic microzones, and steep redox gradients, that select for specialized taxa capable of methane oxidation, sulphate reduction, and hydrocarbon degradation. These naturally occurring habitats may be viewed as the evolutionary and functional foundation for the microbial processes observed at industrially disturbed sediments. The capacity of microbial communities to detoxify hydrocarbons and transform complex carbon substrates represents an ecosystem service that is both economically and ecologically significant, contributing to the natural attenuation and biogeochemical regulation of hydrocarbon-rich seabed environments [54]. By regulating processes such as pollutant detoxification and thereby reducing impacts on meiofaunal and macrofaunal communities, microbes play a critical role in enabling the reestablishment of sensitive species following disturbance [55–57].

### Explanatory power varies across approaches and feature resolution

All approaches consistently resolved impact gradients and identified THC and Ba as the primary driver of community dissimilarity, supporting the use of prokaryotes as sensitive indicators of local impacts from Oil & Gas installations. Shotgun metagenomics provided the highest explanatory power for distinguishing impacted from non-impacted sediments, whereas metabarcoding achieved comparable performance only at the taxonomic level. This indicates that direct access to genomic functional potential offers the strongest signal of environmental disturbance. Metatranscriptomics showed lowest discriminative power although rendering the clearest functional profiles. While metatranscriptomics recovered fewer KOs and ES-relevant processes, these produced clearer functional signatures of environmental impact, suggesting that the expressed fraction captures the most biologically and ecologically meaningful responses. The lower statistical power of metatranscriptomics is primarily attributable to its substantially higher within-group variability, as reflected by consistently higher residual variance and dispersion across KOs, families, and ES-relevant processes. The greater variability among replicates within groups could be explained by methodological biases (RNA degradation, cDNA synthesis errors, removal of ribosomal RNA during library preparation) and the fact that metatranscriptomics captures functional activity, which may fluctuate on short time scales, making it inherently more variable than the static functional potential revealed by metagenomics. By retaining only actively expressed genes and excluding inactive or rare functions, metatranscriptomics reveals functional pathways that are responding to local environmental conditions, therefore resulting in more interpretable functional profiles, even is the underlying statistical discriminatory power is lower.

Taxonomy at family level appeared to be the best feature for distinguishing impacted vs. non-impacted stations across approaches (Table 1), corroborating that this is a well-suited taxonomic rank in the context of benthic environmental status assessment, offering a balance between specificity and sensitivity for prokaryotes [21,58,59]. For metabarcoding data, family community structure discriminated better between impacted and non-impacted sediments than functional profiling, which is not surprising considering that Tax4Fun2 does little more than transforming the underlying metabarcoding data to a new feature space. However, family rank taxonomy performed similarly (PERMANOVA) or, according to dbRDA, even better than functional community structure, *i.e.* using KOs or ESs, also for metagenomics, underscoring a pattern that was not anticipated. In ecosystems with low functional redundancy, shifts in taxonomic composition may translate directly into shifts in functional potential, as the loss or replacement of a few families can remove key metabolic capabilities. This dynamic provides a plausible explanation for why both family-level community structure and KEGG ortholog profiles were similarly effective at distinguishing impacted from non-impacted sites [60,61].

### Functional prediction from 16S metabarcodes sometimes contradicts results from metagenomics

Although 16S-based functional prediction distinguished impacted stations from non-impacted ones and provided an ES-process inventory largely overlapping with that from metagenomics (69%), our study showed little agreement between SIMPER-identified ES-processes between metabarcoding and metagenomics (29% and 26% overlap for ES-processes contributing to impacted and non-impacted communities, respectively, Fig. 4). We found contradicting results for shared ES processes such as methanogenesis from amines, methanotrophy, denitrification, which had relative abundance patterns that did not match between the two approaches. To illustrate the potential significance of this, we considered methanogenesis from amines, which is primarily associated with two archaeal families Methanosarcinaceae and Methanomassiliicoccaceae. Both were detected in the metagenomic and metatranscriptomic datasets but were largely absent from the metabarcoding inventory, likely due to PCR primer bias. As methanogenesis is a key microbial process linked to greenhouse gas emissions, differences in methanogenic potential across impact gradients may have important implications for ecosystem functioning and environmental assessment. This underscores the value of metagenomics for capturing functionally important microbial taxa that may be missed by PCR-based metabarcoding.

Since 16S-based functional prediction tools rely on the assumption that 16S rRNA gene similarity is a reliable proxy for overall genome similarity, discrepancies may also arise when this is violated [62,63]. Our findings highlight that inferring functional potential based on taxonomic profiles may not fully capture the true functional diversity present in complex environmental samples. While Tax4Fun2 can detect specialized metabolic functions (e.g., methane metabolism), its reliance on reference genomes limits its accuracy in complex or poorly characterized environments (and PICRUSt2’s hidden state prediction approach may perform better in this case). Thus, while Tax4Fun2 may serve as a viable tool in North Sea coastal and shelf sediments, its applicability in less well characterized environments may become severely limited. For future applications, metabarcoding remains a valuable tool for environmental impact assessment when restricted to taxonomic composition or SV-level analyses. However, the use of predicted functional profiles derived from 16S data should be approached with caution, as they lack the resolution and reliability needed for robust ecological inference. A promising strategy involves generating metagenomes from a representative subset of samples to establish environmentally relevant, annotated reference genomes. These can then serve as a foundation for inferring functional profiles from SVs in broader metabarcoding datasets. The rationale for incorporating functional data into impact assessments lies in its potential to be less sensitive to geographic variation, and this assumption requires empirical validation through direct metagenomic sequencing.

## Conclusion

Overall, this study showed clear effects of impact of Oil & Gas operations at taxonomy and functional levels regardless of the genomic approach used (metabarcoding, metagenomics or metatranscriptomics). These effects include a shift towards anaerobic, hydrocarbon-degrading, and sulphur-respiring microbial communities. Future studies that incorporate more stations, also at intermediate levels of impact, could provide higher statistical power and enhance the understanding of ecosystem service processes along finer disturbance gradients. The 16S-based function prediction appeared to work reasonably well for impact assessment, while it proved to be problematic for ecological interpretation. Metabarcoding therefore may not be suitable for ES-based monitoring due to the combined limitations posed by primer bias, insufficient genomic data, and prediction approaches. More genomes and stricter algorithms are required for metabarcoding-based functional prediction to become applicable in more diverse and less explored habitats. Our results demonstrate that prokaryotic functional profiles are highly effective for distinguishing impacted sites from non-impacted ones and provide a strong framework for integrating taxonomic and functional assessments of environmental impacts. Metagenomics provided greater statistical discrimination because functional potential is relatively stable across replicates, whereas the inherently dynamic nature of gene expression resulted in greater variability and slightly lower discriminatory power for metatranscriptomics. Nevertheless, by capturing only actively expressed functions, metatranscriptomics produced clearer signatures of pathways responding to environmental impact, facilitating ecological interpretation.

## Acknowledgements

We would like to thank Akvaplan-niva, and the crew of the Ocean Response supply ship, who greatly aided the field work for this study. We thank Rolf Christian Sundt and Linn Pedersen Hocking from Equinor AS for sampling logistics. We thank Sigrid Mugu for valuable input on lab processing. We also thank Anita Skarstad and Ane Kjølhamar from Equinor AS and Thomas Merzi from TotalEnergies AS for project input and support.

## Author contributions

Anders Lanzén (Conceptualization, Validation, Writing—review & editing), Jon Thomassen Hestetun (Conceptualization, Funding acquisition, Writing—review & editing), Thomas G. Dahlgren (Funding acquisition, Supervision, Writing—review & editing), Aud Larsen (Resources, Writing—review & editing), Miriam I. Brandt (Conceptualization, Investigation, Data curation, Project Administration, Writing—original draft), and Andrea Bagi (Conceptualization, Methodology, Data curation, Formal analysis, Writing—original draft, Writing—review & editing).

## Data availability

The raw Illumina sequence data of metagenomic, metatranscriptomic and metabarcoding data have been deposited in the sequence read archive at in the European Nucleotide Archive (ENA) at EMBL-EBI. The metagenomics and metatranscriptomics data for this study have been deposited under accession number PRJEB105097 (https://www.ebi.ac.uk/ena/browser/view/PRJEB105097). Demultiplexed metabarcoding data was submitted under accession number PRJEB109298 (sample accession ERS28573882-ERS28573887 and ERS28573896-ERS28573898). All remaining data generated or analyzed during this study are included in this article and its supplementary information files. The scripts and data require running the scripts for all ecological and statistical analyses are available in the following Zenodo repository: https://doi.org/10.5281/zenodo.18301504.

