## Supplementary figures and images for "Taxonomic *vs*. Functional Diversity for the Impact Assessment of Offshore Oil & Gas Activities – An Exploratory Study on Benthic Prokaryotes"

### Supplementary Figure S1

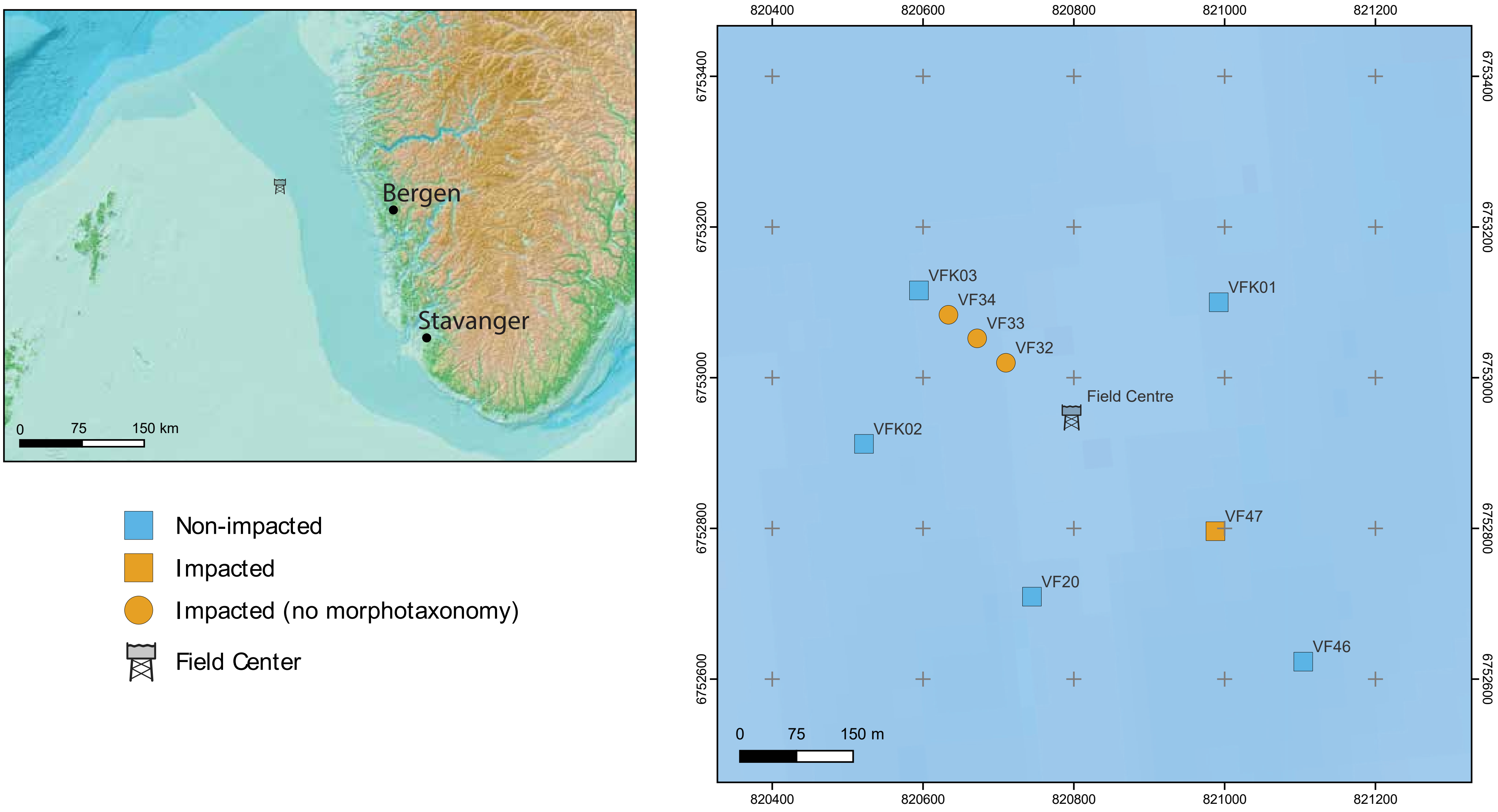

### Supplementary Figure S2

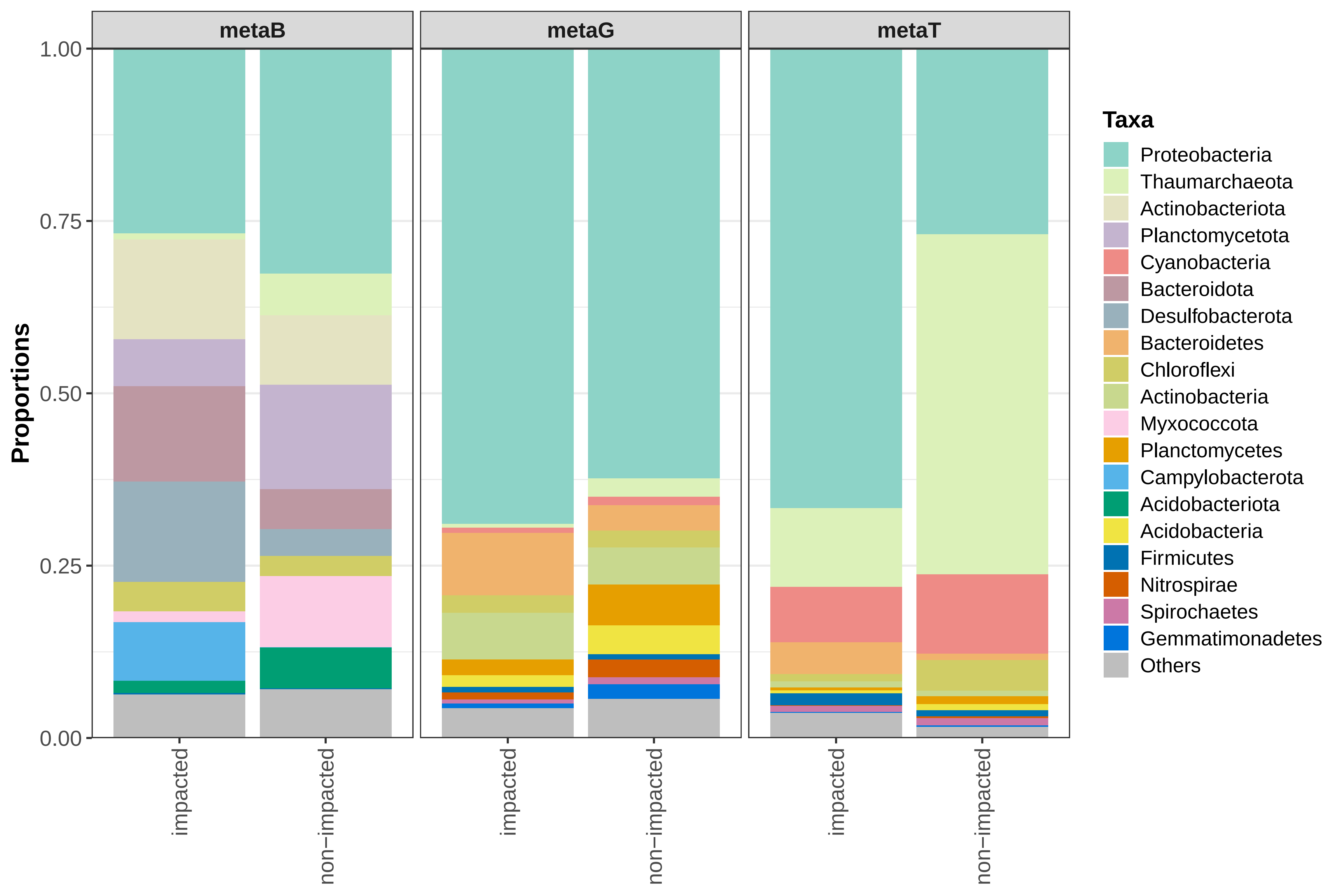

### Supplementary Figure S3

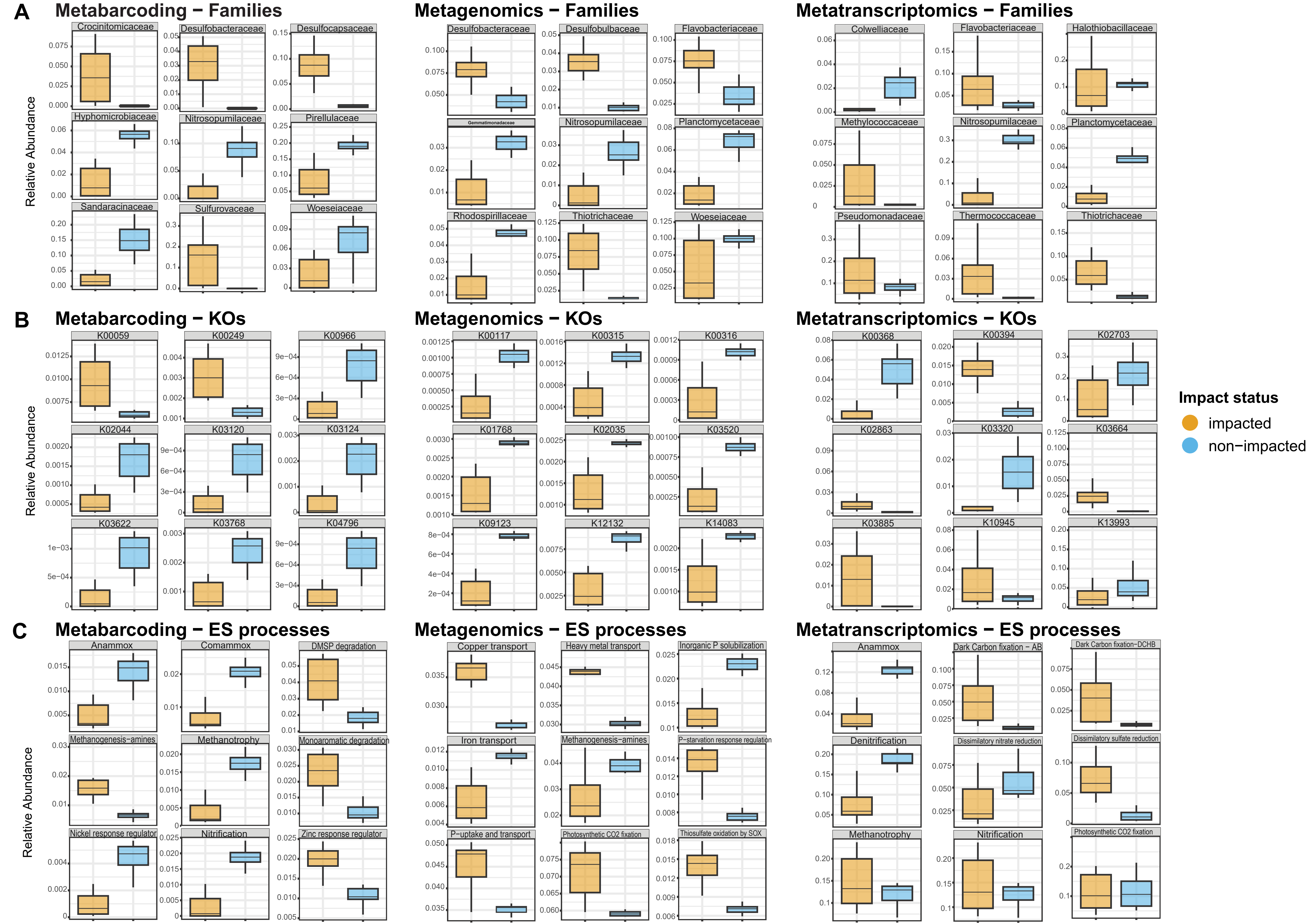

### Supplementary Figure S4

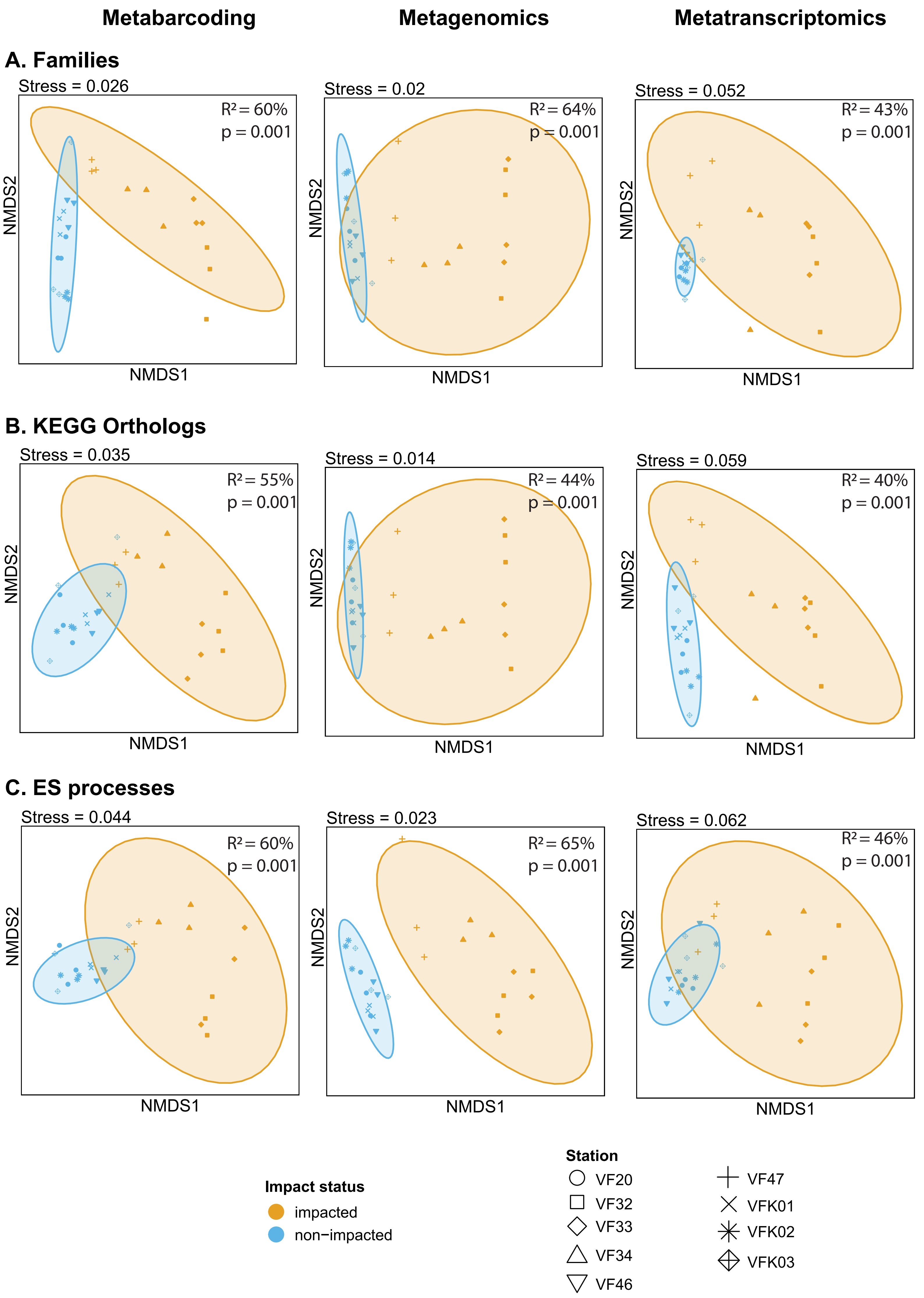
