## Supplementary Methods for "Taxonomic *vs*. Functional Diversity for the Impact Assessment of Offshore Oil & Gas Activities – An Exploratory Study on Benthic Prokaryotes"

² AZTI, Marine Research, Basque Research and Technology Alliance (BRTA), Pasaia, Spain

^3^ IKERBASQUE, Basque Foundation for Science, Bilbao, Spain

^4^ Department of Marine Sciences, University of Gothenburg, Gothenburg, Sweden

### Supplementary Materials and Methods

##### Site Description and Sample Collection

Sediment was collected on the Norwegian continental shelf around the Veslefrikk offshore installation, which has been operational since 1989 and was decommissioned in 2022 (Fig. S1, Table S1). Sampling was performed during the routine Norwegian offshore monitoring program [1], by Akvaplan-niva in May 2022 (full reports available at [2]), with measurement of both physicochemical parameters and macrofauna biotic indices. Sediment was collected at 9 stations (5 non-impacted, 4 impacted, depth range 175-178 m) with five double-chamber Van Veen grabs (surface area 0.15 m^2^) per station, three of those also subsampled for chemistry. Sediment THC content of surface sediments (0-1 cm) was measured via gas chromatography (GC/FID) and heavy metals (As, Ba, Cd, Cu, Cr, Hg, Pb, Zn) were quantified via ICP-SFMS, following ISO standards 17294-1,2. Chemical profiles of the 9 sampled stations (Fig. S1) showed that THC values ranged from 210 to 27,948 mg/kg (13,269±11,791 mg/kg) and from 23.7 to 64.7 mg/kg (37.5±14.9 mg/kg) for impacted and non-impacted stations, respectively. Barium concentrations ranged from 6,517 to 12,067 mg/kg (8,727±2,209 mg/kg) and from 680 to 4,570 mg/kg (2,961±1,627 mg/kg) for impacted and non-impacted stations, respectively (Table S1). Samples from three stations (VF32, VF33, VF34) were not processed for granulometry and morphotaxonomy due to their high hydrocarbon content (THC range 4,000 - 30,000 mg/kg) [3] . Signs of disturbance, based on macrofauna assessment, were observed only at VF47 (VFR05) among the stations sampled for macrofauna analysis.

Sediment for eDNA and eRNA was taken from the top 2 cm from three locations within each “chemistry” grab, either from the full grab chamber (stations where morphotaxonomy was not carried out) or from the small 0.05 m^2^ compartment (stations where morphotaxonomy was carried out). Gloves and surgical masks were worn when sampling, and a new disposable spatula was used for each grab to reduce contamination risks. For metagenomics and metatranscriptomics, sediment was pooled in a 10 ml cryotube, giving around 10 g for each grab sample. The cryotubes were flash-frozen in liquid nitrogen, put into a small 80x120 mm zip lock bag, and stored at -20°C immediately after sampling. A sampling blank was also collected using an empty cryovial opened during sampling and put into its own zip lock bag. For eDNA metabarcoding, the process was equivalent, except that sediment was pooled in a 50 ml polypropylene tube for each grab, yielding around 30 g. Each 50 mL tube was then stored at -20°C on board. All samples were shipped at -20°C to the laboratory in Bergen, Norway, and then transferred to a -80°C freezer.

##### Nucleic acid extractions

Work was performed in an over-pressured cleanroom wearing appropriate equipment for low-DNA samples (coat, mask, hairnet, disposable gloves). All work surfaces were washed with fresh 10% bleach solution and rinsed with ultrafiltered water at the beginning and end of each extraction round.

###### Standard eDNA extraction.

Sediment was left to thaw in its tubes overnight at 4°C. Sediment was homogenized manually with a single-use spatula for 1 min and ~0.5 g of sediment was weighed into triplicate pre-labelled PowerBead tubes (ref 19301, as in the DNeasy PowerSoil Pro kit). A subsampling blank, i.e. an empty PowerBead tube, was opened during subsampling. Sediment in triplicate PowerBead tubes was stored at -20ºC before extraction using a semi-automated protocol combining PowerBead tubes and CD1 solution (ref 47016, Qiagen, Hilden, Germany), a Precellys 24 homogenizer (Bertin Technologies, Montigny-le-Bretonneux, France) and a Qiagen QIAsymphony SP instrument (Qiagen, Hilden, Germany). Briefly, sediment in PowerBead tubes was thawed on ice, spinned down and 800 µl of CD1 solution was added to the tubes. Cell lysis was then performed on a Precellys 24 instrument with a medium intensity program (4000 rpm for 1 min), following recommendations in [4]. After centrifugation (15,000 x g, 1 min, room temperature), the supernatant was transferred to a labelled and UV-sterilised 2 mL Sarstedt microtube (Sarstedt, Nümbrecht, Germany) and loaded to a QIAsymphony SP instrument for extraction using the QIAsymphony DSP DNA Mini Kit (Qiagen, Hilden, Germany) and the Tissue_LC_200_V7_DSP (low content) protocol with a 100 µl elution volume. Extraction negative controls were included in each session. DNA concentrations were measured with the dsDNA HS Assay Kit (5 µl extract) on a Qubit 3.0 fluorometer (ThermoFisher Scientific).

###### Joint DNA/RNA extractions.

Sediment in cryotubes was left to thaw one hour on ice. Joint RNA/DNA extractions were performed with fresh phenol/chloroform/isoamyl alcohol solution (pH 6.5-8.0, 25:24:1) using the RNeasy PowerSoil Kit (12866-25) combined with the accessory DNA elution kit (12867-25, Qiagen, Hilden, Germany). About 5 g of sediment were used, following the manufacturer’s suggestions for wet sediments. Extraction controls were performed alongside sample extractions. All RNA extracts were DNase-treated with the TURBO-DNA-free Kit (AM1907, ThermoFisher Scientific, Waltham, MA, USA) in 50 µL reactions to remove any contaminant DNA. Concentrations in final extracts were measured using a Qubit 3.0 fluorometer with the RNA HS and the dsDNA HS Assay Kits (ThermoFisher Scientific).

##### Library Preparation and Sequencing

Jointly extracted eDNA and eRNA were sent to Novogene Europe (Oxford, United Kingdom) for library preparation and sequencing and processed using Novogene’s in-house protocols. Genomic DNA was randomly sheared into short fragments, which were end repaired, A-tailed and further ligated with Illumina adapter. The fragments with adapters were PCR amplified, size selected, and purified and sequenced on an Illumina Novaseq 6000 platform. The libraries consisted of fragments with 350 bp nominal insert size. Paired end sequencing was performed with 300 cycles, resulting in 150 nucleotide long reads for each read pair. The anticipated sequencing depth was 40 Gb (corresponding to 130 million reads per sample). Metatranscriptomic libraries were prepared following ribosomal RNA depletion step (removal of rRNA by using oligos complementary to rRNAs). Like for the metagenomic libraries, PE150 strategy was used, however, a sequencing depth of 20 Gb (corresponding to 65 million reads) was chosen here. The metagenomics and metatranscriptomics data for this study have been deposited in the European Nucleotide Archive (ENA) at EMBL-EBI under accession number PRJEB105097 (<https://www.ebi.ac.uk/ena/browser/view/PRJEB105097>).

For metabarcoding, the prokaryote 16SV4V5 region was amplified from the standard eDNA extracts with the 515F-Y/926R primer pair [5], using primers with 8 nucleotides (nt) sample tags on the 5’ end, enabling nested multiplexing of PCR products as described in [6]. The triplicate extracts were pooled for PCR, as recommended by [7] and single PCR amplifications were carried out per sample, as the study aim was to resolve the core community structures [8]. The PCR reactions (25 μL final volume) contained 10 ng or less of DNA template with 0.3 μM final concentration of each primer, 0.2 mg/mL of BSA, and 1X KAPA plant PCR Buffer (KK7252, Roche, Basel, Switzerland). PCR thermocycling conditions were 95 °C for 3 min; then 30 cycles of 20 s at 95 °C, 30 s at 50°C, 45 s at 72 °C; followed by 72 °C for 5 min. PCR-negative controls were performed along samples PCRs, and a mock community was included as positive control (ZYMObiomics microbial community DNA standard, ref D6306, ZYMO research, Irvine, CA, USA). PCR products were quantified by high-resolution capillary electrophoresis using a QIAxcel Instrument (Qiagen, Hilden, Germany). Equimolar pools of raw PCR products were prepared by pooling ~50 ng per PCR product, to ensure balanced representation of each sample in the sequencing run. The libraries were then purified using KAPA Pure Beads at a 1.0X ratio (Roche, Basel, Switzerland). Amplicon libraries were sent to Novogene Europe for PCR-free library preparation using the NEBNext® Ultra™ II DNA Library Prep Kit (Cat No. E7645, New England Biolabs, Ipswich, MA, USA) with alterations: Y shaped adaptors were ligated to the amplicons (replacing the U excision), and one cycle of PCR was performed to blunt the ends of the adaptors. After this, the PCR-enrichment step in the kit protocol was entirely skipped and the library was sent for quality control. Libraries were quantified via Qubit and real-time PCR, and fragment size distribution was verified on a high-throughput microfluidic capillary electrophoresis system (Bioanalyzer, Agilent, Santa Clara, CA, US) to calculate molarity. Quantified libraries (1.5-30 nmol/L) were pooled according to effective library concentration, adjusting volumes based on data amount required (4 Gb, here ~200,000 reads per sample). Final pooled libraries were sequenced on Novaseq 6000 instruments (Sequencing System Guide 1000000019358 v17, Illumina, San Diego, CA, USA) in 250 base pairs paired-end mode, with final loading concentrations of 300-600 pM and a 40% PhiX spike-in (NovaSeq 6000 Denature and Dilute Guide document #1000000106351) using NovaSeq 6000 SP Reagent Kits. Raw data was demultiplexed at Novogene based on the index in the sequencing adapters. Adapters were removed, and raw data filtered by removing reads containing adapter sequences, removing reads with N > 10% and removing reads containing bases with quality scores <= 5 over 50% of the read.

##### Sequence Processing and Annotation

*Metagenomics*: Low quality bases (Q-score ≤ 38 over 40 bp), reads containing more than 10 undetermined (“N”) nucleotides and reads which overlapped with adapter sequences (over 15 bp) were removed during the quality control step (Readfq v8). MEGAHIT (v1.0.4-beta) with k-mer size of 55 was then used for *de novo* assembly of quality filtered and trimmed reads separately for each sample at first. The scaffolds were cut off at N to generate continuous sequences within scaffolds without N (named scaftigs). Reads were mapped back onto the scaftigs with SoapAligner (v2.21) and a subsequent mixed assembly was performed on all unutilized (unmapped) reads. Scaftigs shorter than 500 bp were removed prior to gene prediction. Open reading frames (ORFs) were identified with MetaGeneMark (v3.05). Only ORFs longer than 100 nt were kept and subsequently dereplicated with CD-HIT (v4.5.8) using 95% identity cutoff and minimum coverage of 90% (parameters: -c 0.95, -G 0, -aS 0.9, -g 1, -d 0). The longest sequences were chosen as representative “unigenes” (unique gene sequences). Reads were mapped onto the unigenes using SoapAligner (v2.21) (parameters: -m 200, -x 400, identity = 95%) and length-normalized counts (G_k_) were calculated. The formula for calculating G_k_ values is equivalent to the calculation of transcripts per million (TPM) values without the multiplication step (*10^6^). G_k_ values sum to 1 on a per sample bases, instead of 1 million (as for TPM).

*Metatranscriptomics*: Reads that contained adapter sequences, over 10 % of N nucleotides and having Q-scores ≤ 20 over 50% of the read length were removed using fastp (v0.23.1) Ribosomal and transfer RNA sequences were removed by mapping to the SILVA and NCBI databases. Filtered sequences were then assembled using Trinity (v2.4.0), and assembled transcripts were dereplicated with CD-HIT (v4.6). Due to computational limitations, it was not possible to utilize reads from all samples combined in the assembly process. Instead, only reads from 9 samples, *i.e.*, from one replicate per station (VFK1_1, VFR04_1, VFR20_1, VFR05_1, VFR32_1, VFR33_1, VFK2_1, VFK3_1, VFR34_1) were used to create an assembly. The longest transcript sequence (called unigene) was kept as representative of each transcript cluster. The alignment-based tool, RSEM (v1.2.15) with bowtie alignment (mismatch 0) was used to establish the fragments per kilobase per million (FPKM) counts of the transcripts.

For both metagenomics and metatranscriptomics, unigenes were aligned to Novogene’s microNR database (includes sequences of Bacteria, Fungi, Archaea and Viruses extracted from NCBI’s NR database, version 2018-01-02) using DIAMOND v0.9.9 (blastp, e-value ≤ 1e-5, -more-sensitive) and taxonomy was assigned using MEGAN, lowest common ancestor (LCA) algorithm. The abundance of a species is calculated from the sum of abundances of the genes annotated to that species. For functional annotation, unigenes were mapped to Kyoto Encyclopedia of Genes and Genomes (KEGG, version 2018-01-01) using DIAMOND v0.9.9 (blastp, e-value 1e-5, -more-sensitive) and best BLAST hit was selected.

*Metabarcoding:* data demultiplexing and processing was carried out using the scripts *01_demultiplex.sh,* and *02_runDADA2-v2.R* available from https://github.com/lanzen/MetaBridge*.* In summary, *cutadapt* v.4.5 [9] was used to demultiplex samples based on the 8 nt sample tags, and then to re-orient reads and remove primers, discarding reads with incorrect or incomplete primer sequences. Demultiplexed data was submitted to ENA under accession number PRJEB109298 (sample accession ERS28573882- ERS28573887 and ERS28573896- ERS28573898). Pairs of Illumina reads were then denoised using DADA2 v.1.28 (truncLen = 220, minOverlap = 20, maxOverlapMismatch = 1, lengthMin=300, lengthMax=420) with a modified Loess function to account for Novaseq sequencing (see https://github.com/benjjneb/dada2/issues/1307), resulting in a set of unique amplicon sequence variants (SVs) and SV distribution tables. Finally, taxonomic assignment was conducted using CREST v4.3.7 [10] (https://github.com/xapple/crest4), with the SSUOME1.1 database (combining SILVA SSURef v138 and EUKARYOME v1.9.3). ASV tables were refined in R v.4.4 [11]. Potential contaminants were removed based on subsampling, extraction and PCR blanks using the *decontam* package v.1.24.0 [12], with the combined frequency and prevalence-based methods. Contaminant identification was performed independently for each library, using the *batch* option. Cross-contamination was then reduced using a custom R function based on the UNCROSS algorithm [13], changing sample-specific OTU abundances to zero when their relative abundance in a sample was < 1% of their average abundance across samples. Sequences matching known mock species from sequenced mock community samples were then used to estimate the degree of remaining cross-contaminants, after which all such sequences were eliminated. Finally, all OTUs unclassified at phylum rank or assigned to eukaryotic organelles were removed.

Functional prediction was performed using Tax4Fun2 (v1.1.5) using the refined ASV table and associated sequences [14] using SILVA Ref99NR as taxonomic reference database and a minimum identity to reference sequences of 97% and *normalize_pathways* set to FALSE, following the manual [15]. The output of tax4fun2 is a relative abundance table with KEGG Ortholog numbers as features.

##### Ecosystem Services Relevant Functions

To enable comparisons based on functions relevant to ecosystem services (ES), a curated list of KEGG Orthologs (KOs) was compiled using KEGG and published sources (Table S2). The selection focused on two broad ES categories: (1) supporting services like nutrient and element cycling, and (2) regulating services such as detoxification of hydrocarbons and heavy metals. Where KEGG modules existed (e.g., nitrogen cycling: M00531, M00529), they were used to link ES functions to KOs. For processes lacking modules (e.g., organic phosphorus mineralization, alkane degradation), essential genes and their KO identifiers were manually gathered. Phosphorus-related KOs were sourced from [16], alkane degradation KOs focused on key oxygenases, and metal-related genes were drawn from recent studies [17,18]. Carbon cycling KOs were selected for their roles in CO₂ fixation and methane dynamics, reflecting their impact on greenhouse gas fluxes.

##### Ecological Data Analysis and Statistics

Prevalence filtering was performed at feature level (ORFs, transcripts and ASVs for metagenomics, metatranscriptomics and metabarcoding, respectively) to retain only features that were present in all 3 grab replicates at each station.

For all the statistical tests, performed in R (v4.4.0), Bray-Curtis distances calculated from Hellinger transformed data using vegan (v2.6.10), were used to quantify community dissimilarities.

Non-Metric Dimensional Scaling (NMDS) was performed using metaMDS and visualized using ggplot2 (v3.5.1). Permutational Multivariate Analysis of Variance (PERMANOVA, adonis2, 999 permutations) was used to test the effect of impact on prokaryote communities. Multivariate homogeneity of groups dispersions was evaluated with betadisper function (999 permutations).

For each approach, bar plots of mean relative abundances across impact categories showing compositional structure at family, KEGG Ortholog and ES-relevant metabolic process level were created using ggplot2 (v3.5.1).
